# Host-pathogen coevolution increases genetic variation in susceptibility to infection

**DOI:** 10.1101/474924

**Authors:** Elizabeth ML Duxbury, Jonathan P Day, Davide Maria Vespasiani, Yannik Thüringer, Ignacio Tolosana, Sophia CL Smith, Lucia Tagliaferri, Altug Kamacioglu, Imogen Lindsley, Luca Love, Robert L Unckless, Francis M Jiggins, Ben Longdon

**Author notes:** joint first authors (Elizabeth ML Duxbury, Jonathan P Day). equal contributions, joint senior and corresponding authors, and (Francis M Jiggins, Ben Longdon).

## Abstract

It is common to find considerable genetic variation in susceptibility to infection in natural populations. We have investigated whether natural selection increases this variation by testing whether host populations show more genetic variation in susceptibility to pathogens that they naturally encounter than novel pathogens. In a large cross-infection experiment involving four species of *Drosophila* and four host-specific viruses, we always found greater genetic variation in susceptibility to viruses that had coevolved with their host. We went on to examine the genetic architecture of resistance in one host species, finding that there are more major-effect genetic variants in coevolved host-parasite interactions. We conclude that selection by pathogens increases genetic variation in host susceptibility, and much of this effect is caused by the occurrence of major-effect resistance polymorphisms within populations.

## Introduction

From bacteria to plants and insects to humans, it is common to find considerable genetic variation in susceptibility to infection in natural populations [1-4]. This variation in susceptibility can determine the impact of disease on health and economic output [5-8]. In nature and breeding programs, it determines the ability of populations to evolve resistance to infection. Insect populations, like those of other organisms, typically contain considerable genetic variation in susceptibility to infection [2, 4, 9, 10], and provide a convenient laboratory model in which to investigate basic questions about how this variation is maintained [11]. Within vector species like mosquitoes, resistant genotypes are less likely to transmit parasites, and this has the potential to reduce disease in vertebrate populations [12]. Where pathogens are contributing the decline of beneficial species like pollinators, high levels of genetic variation may allow populations to recover [13]. Understanding the origins of genetic variation in susceptibility is therefore a fundamental question in infectious disease biology.

As pathogens are harmful, natural selection is expected to favour resistant host genotypes. Directional selection on standing genetic variation will drive alleles to fixation, removing variants from the population [14-16]. However, as directional selection also increases the frequency of mutations that change the trait in the direction of selection, at equilibrium it is expected to have no effect on levels of standing genetic variation (relative to mutation-drift balance; [17]). However, selection mediated by pathogens may be different. Coevolution with pathogens can result in the maintenance of both resistant and susceptible alleles by negative frequency dependent selection [18, 19]. Similarly, when infection prevalence exhibits geographical or temporal variation, selection can maintain genetic variation, especially if pleiotropic costs to resistance provide an advantage to susceptible individuals when infection is rare [20-22]. Even when there is simple directional selection on alleles that increase resistance, the direction of selection by pathogens may frequently change so populations may not be at equilibrium. If selection favours rare alleles – such as new mutations – directional selection can transiently increase genetic variation during their spread through the population [23-25].

To examine how selection by pathogens affects levels of genetic variation we used a natural host-parasite system; *Drosophila* and sigma viruses [26, 27]. Sigma viruses are a clade of insect RNA viruses with negative-sense genomes in the family Rhabdoviridae [28-31]. They are vertically transmitted, and each sigma virus infects a single host species, simplifying studies of coevolution [28, 31]. In *Drosophila melanogaster* infection reduces host fitness by approximately 25% and prevalence in wild populations is typically around 10% [32, 33]. As such, there is the necessary selective pressure for resistance to evolve [25, 34]. In *D. melanogaster*, three major-effect resistance alleles have been identified [2, 11, 24, 25, 34-36]. There has been a recent sweep of genotypes of *D. melanogaster* sigma virus (DMelSV) that are able to overcome one of these host resistance genes [37-39]. Given the power of *Drosophila* genetics, this system is an excellent model of a coevolutionary arms race between hosts and parasites.

Sigma viruses offer a novel way to test how coevolution with a pathogen alters the amount of genetic variation in host susceptibility. As sigma viruses are vertically transmitted, we can be certain of which hosts and viruses are natural coevolving combinations, and which do not naturally coevolve. In this study we have used four species of *Drosophila* (*D. affinis, D. immigrans, D. melanogaster* and *D. obscura*) that shared a common ancestor approximately 40 million years ago [40, 41], and their natural sigma viruses (DAffSV, DImmSV, DMelSV and DObsSV respectively) [29, 30, 42]. We have tested whether selection by parasites increases the amount of genetic variation in host susceptibility by comparing viruses that naturally infect and coevolve with each species to non-coevolving viruses. We then examined how selection by pathogens alters the genetic architecture of resistance by mapping loci that confer resistance to DAffSV, DMelSV and DObsSV in *D. melanogaster*.

## Results

### Genetic variation in susceptibility to infection is greatest in coevolved host-virus associations

To test whether selection by parasites increases genetic variation in susceptibility to infection, we compared coevolved host-virus associations with novel associations that have no history of coevolution. We used four different species of *Drosophila*, each of which is naturally host to a different sigma virus [26-28, 30, 31]. We collected four species from the wild and created genetically diverse populations in the laboratory. Using flies from these populations we crossed single males to single females to create full-sib families. The progeny of these crosses were then injected with either the virus isolated from that species, or a virus isolated from one of the other species (Figure 1). Studies on DMelSV have shown that those loci providing resistance against naturally acquired infections in the lab and wild also provide resistance when the virus is injected, and transmission rates are correlated to viral load [2, 27, 43-46]. Fifteen days post infection we extracted RNA from the flies and measured viral load by quantitative RT-PCR. This allowed us to estimate the genetic variance (*V_G_*) in viral load from the across-family variance. In total we infected 52,592 flies and measured the viral load in 4295 biological replicates across 1436 full-sib families (details of sample sizes in Supplementary Methods).

**Figure 1.**
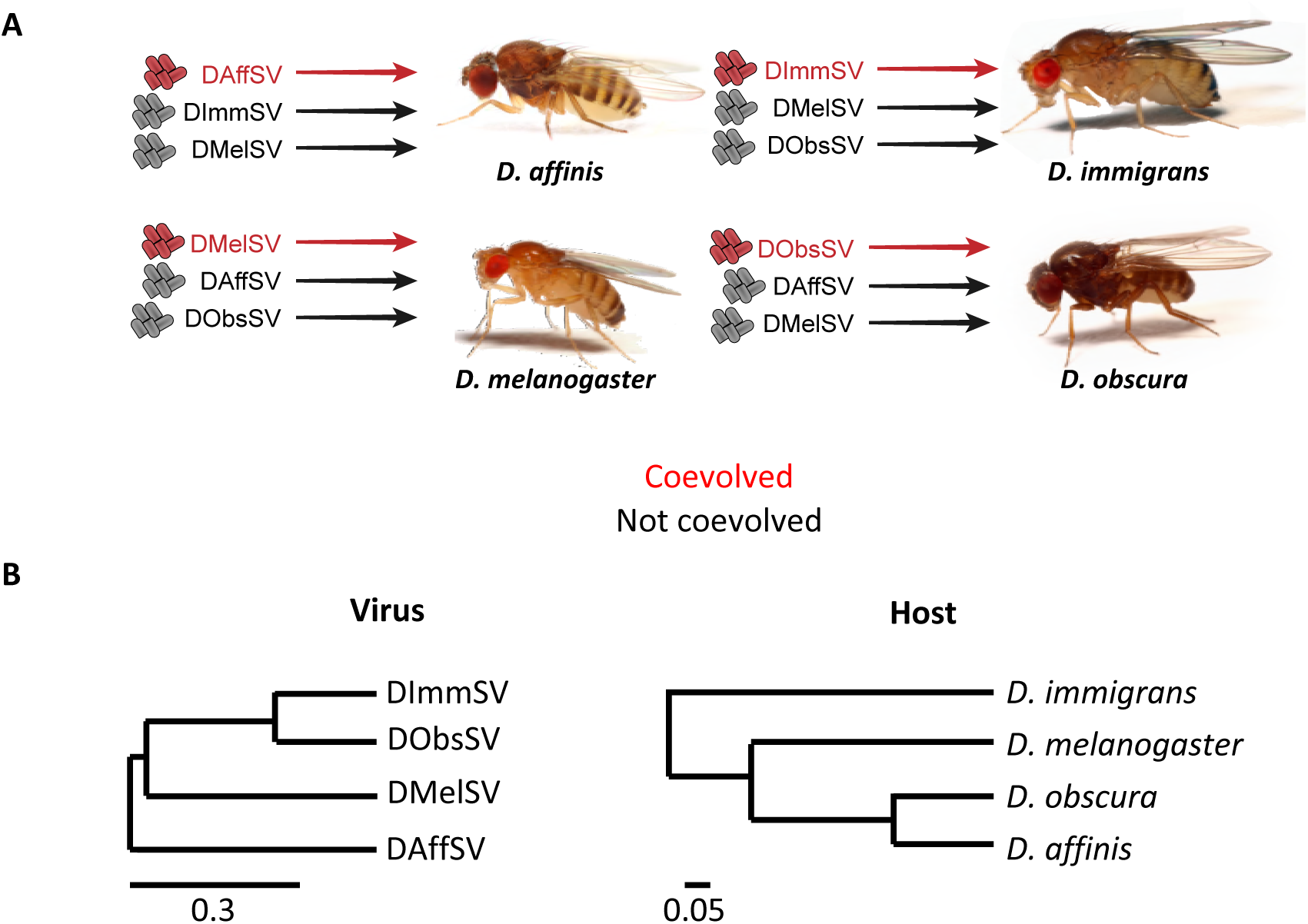
Experimental design and phylogenies of host and parasite. A) Four species of *Drosophila* were independently infected both with a sigma virus with which they have coevolved in nature (red) and two viruses that naturally infect another species (black). **B)** Phylogenies of the sigma viruses and their *Drosophila* hosts, redrawn from [29, 47]. Scale bars represent substitutions per site under a relaxed clock model. Sigma viruses are highly divergent from one another with amino acid identities of <55% in the most conserved gene. The four species of *Drosophila* are estimated to have shared a common ancestor approximately 40 million years ago [40, 41].

Within populations of all four species, we found significantly greater genetic variance in susceptibility to the sigma virus that naturally infects that species compared to viruses from other species (Figure 2 and Supplementary Tables S1 and S2). These different variances reflect considerable differences in the mean viral load between families (Figure 2A). For example, when families of *D. obscura* in the 2^nd^ and 98^th^ percentile were compared, there was a 1294 fold difference between the viral loads of the coevolved virus in the families (Figure 2A). In contrast, for the non-coevolved viruses there was a 27 fold difference in DMelSV loads and a 19 fold difference in DAffSV loads (see Figure 2 for statistics). In *D. affinis* the equivalent statistics were a 26 fold difference in the load of the coevolved virus, compared to a 7-fold and 3 fold difference in DMelSV loads and DImmSV loads. In *D. immigrans* there was an 11-fold difference for the coevolved virus compared to a 2-fold difference for DObsSV and 3-fold for DMelSV. In *D. melanogaster* there was a 9-fold difference for the coevolved virus loads compared to a 4-fold difference for DObsSV and 5 fold for DAffSV. Across the four species, there was no consistent difference between the mean viral load in coevolved versus non-coevolved associations, and the genetic variance in viral load was not correlated with the mean viral load (Supplementary Figure S1, Spearman’s correlation: *ρ*= −0.38, S=296, *P*=0.22).

**Figure 2.**
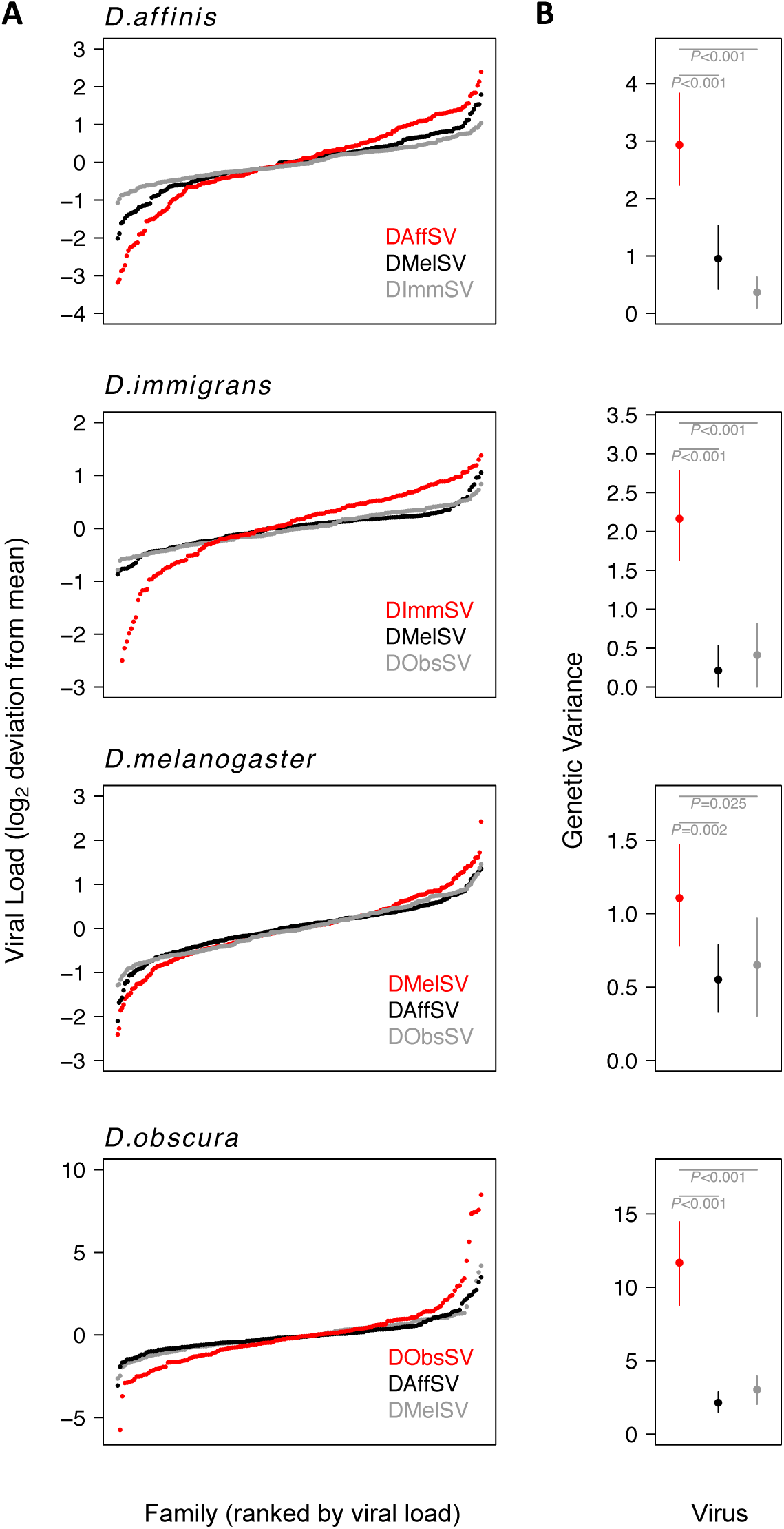
Genetic variation in susceptibility to coevolved and non-coevolved viruses. The viral load was measured 15 days post infection by quantitative RT-PCR relative to a *Drosophila* reference gene (*RpL32*). **A)** The points show model prediction family means from our GLM and are centred on zero. The number of families in each panel was down-sampled so the same number of families is shown for each virus. Coevolved host-virus associations are in red. **B)** The genetic variance in log_2_ viral load was estimated from the between family variance assuming that all genetic variance is additive. The bars are 95% credible intervals. Posterior probabilities for significantly different genetic variances are shown in grey (see Table S1 and S2).

### Major-effect genetic variants that are known to provide resistance to DMelSV do not protect against other viruses

To examine whether the genetic basis of resistance to coevolved and non-coevolved viruses was different we estimated the genetic correlations (*r_g_*) in their viral loads. In *D. melanogaster* these were 0.40 for DMelSV-DAffSV, (95% CIs: 0.20, 0.61) and 0.25 for DMelSV-DObsSV (95% CIs: −0.01,0.47). In the other species our estimates sometimes had wide credible intervals, but the genetic correlations between coevolved and non-coevolved viruses were mostly below 0.5 (Table S3). Therefore if natural selection increases genetic variation in susceptibility to a natural pathogen, there is expected to be a smaller effect on non-coevolved viruses.

In *D. melanogaster* a substantial proportion of the genetic variance in susceptibility to DMelSV is explained by major-effect variants in the genes *CHKov1* and *p62* (also known as *Ref(2)P*) [11, 24, 25, 36, 48]. The resistant allele in each of these genes has arisen recently by mutation and been driven up in frequency by natural selection, presumably due to the presence of DMelSV in natural populations [24, 25]. Therefore, if these genetic variants confer resistance to DMelSV but not the other sigma viruses, then this may explain the differences in genetic variance that we observed.

To examine whether *CHKov1* or *p62* contributed to the differences in genetic variance we observed in *D. melanogaster*, we genotyped the parents of the full sib families for variants that confer resistance (Table S4). Assuming the effects of the resistant alleles are additive, we estimated that the load of the coevolved virus DMelSV was more than halved in homozygous *CHKov1* resistant flies compared to susceptible flies (reduction in log_2_ viral load = 1.2, 95% CI = 0.6, 1.8). In contrast we found no significant effect of this gene on loads of the non-coevolved viruses (DAffSV = −0.2, 95% CI = 0.2, −0.8; DObsSV = −0.4, 95% CI = −0.04, 1.0). The resistant allele of *p62* was present at such a low frequency (1.5%) in the population that we lacked of statistical power to investigate its effects.

To confirm these results we infected 1869 flies from 32 inbred *D. melanogaster* lines [49] that had known *CHKov1* or *p62* genotypes. The effect of these variants was greater on the naturally occurring virus than the viruses from other species (Figure 3; effect genotype on DMelSV load: *F_2,28_*=13.2, *P*=0.00001; DAffSV: *F_2,29_*=4.9, *P*=0.01; DObsSV: *F_2,29_*=5.7, *P*=0.01).

**Figure 3.**
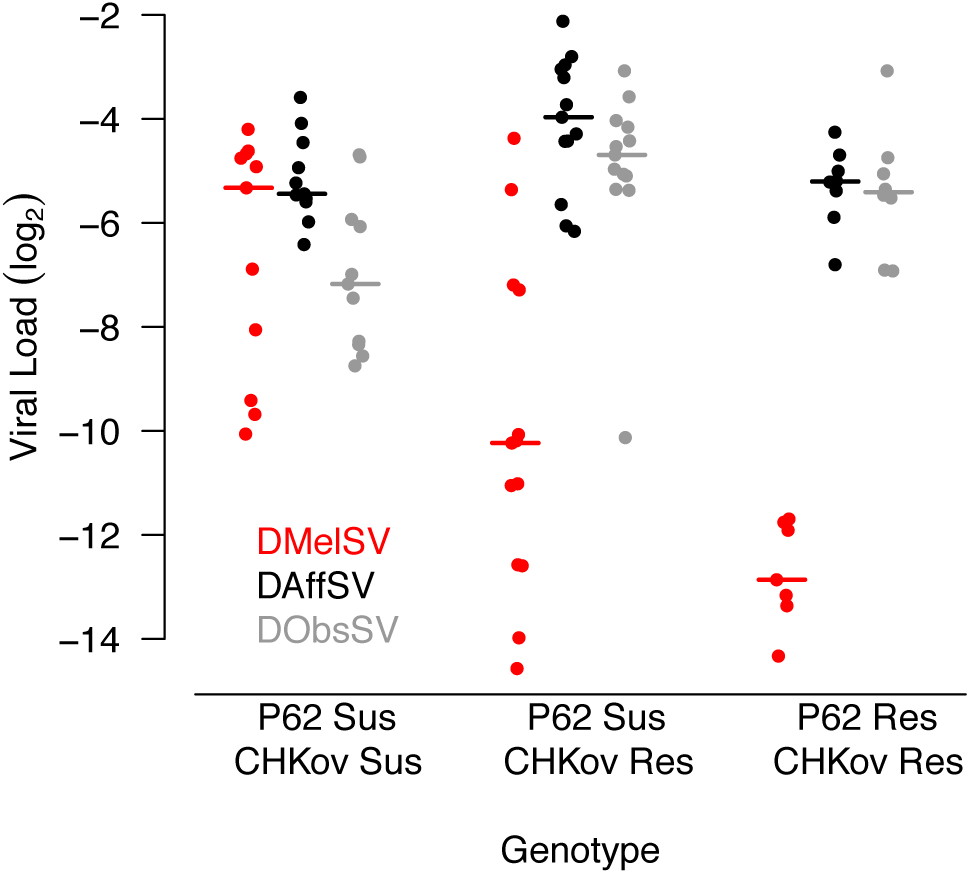
Viral load in *D. melanogaster* lines carrying different alleles of *CHKov1* and *p62*. Each point is the viral load of a separate inbred fly line carrying the resistant (R) or susceptible (S) allele of *P62* or *CHKov1.* Horizontal bars are medians. Viral load was measured 15 days post infection by quantitative RT-PCR relative to a *Drosophila* housekeeping gene (*RpL32*).

### There are a greater number of major-effect variants in coevolved host-virus associations

To investigate how coevolution shapes the genetics of resistance in an unbiased way, we mapped loci controlling resistance using a *D. melanogaster* advanced intercross population (the DSPR panel [50]). This population samples genetic variation in a small number of genotypes from around the world (the experiments above sampled many genotypes from a single location). It was founded by allowing two sets of 8 inbred founder lines to interbreed in large populations for 50 generations, then creating recombinant inbred lines (RILs) whose genomes are a fine-scale mosaic of the original founder genomes. We used 377 RILs from these populations, which have up to 15 alleles of each gene (one founder line is shared between the two populations). We infected 15,916 flies across 1362 biological replicates with DMelSV, DAffSV or DObsSV and measured viral load as above.

We first estimated the genetic variance in viral load within our mapping population. The results recapitulated what we had found above in a natural population of flies — there was considerably more genetic variation in susceptibility to the coevolved virus than the non-coevolved viruses (Figure 4A, filled circles). Therefore, our earlier result from a single population holds when sampling flies from across 6 continents, although the magnitude of the effect is considerably greater in this mapping population.

**Figure 4.**
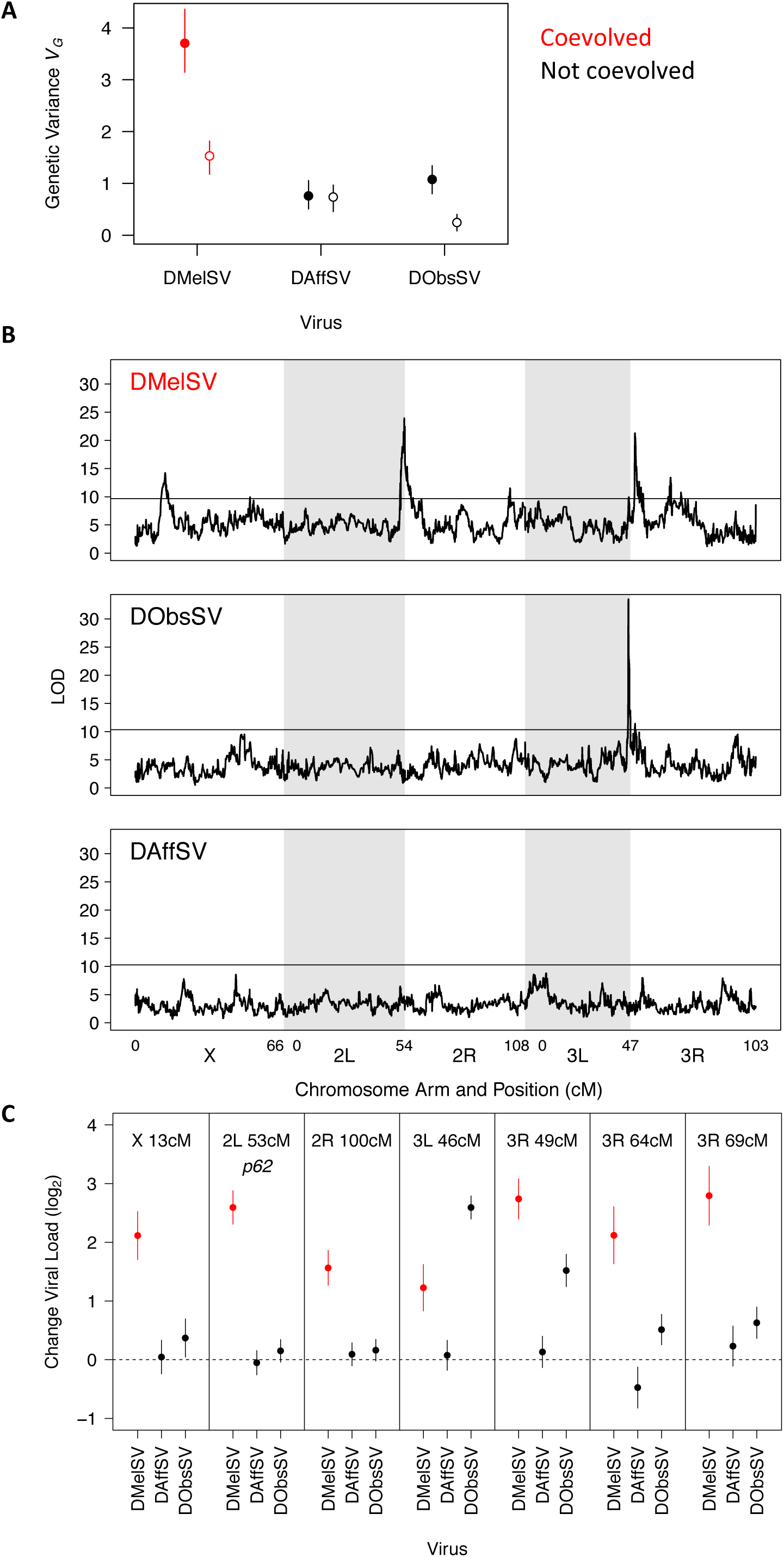
The genetic architecture of resistance to coevolved and non-coevolved viruses in*D. melanogaster*. A) The genetic variance in viral load within the mapping population (filled circles). The open circles are estimates of the genetic variance after accounting to the effects of the QTL in panel C. Error bars are 95% credible intervals. (B) QTL affecting viral load. The horizontal line shows a genome-wide significance threshold of *P*<0.05 that was obtained by permutation. (C) The effect of the seven QTL detected on the load of the three viruses. Only QTL that remained were significant following multiple regression with all the loci are shown. The coevolved virus is shown in red.

To examine the genetic basis of virus resistance, we looked for associations between genotype and viral load across the genome (Figure 4B). In the coevolved association (DMelSV) we identified seven QTL associated with resistance, compared to one that affects DObsSV and none affecting DAffSV (Table S5; this excludes one DMelSV on the X chromosome that did not remain significant after accounting for the other QTL). The QTL affecting DObsSV also has a significant effect on DMelSV. One of the QTL corresponded to *p62* (2L 53cM). The susceptible allele of *CHKov1* was not present in the fly lines assayed.

To examine the effect that the QTL have on viral load, we first split the founder alleles into a resistant class and a susceptible class (see methods) and then estimated the difference in viral load between the functionally distinct alleles. Six of the seven QTL resulted in greater reductions in the load of the coevolved virus (DMelSV) than the viruses isolated from other species (Figure 4C). There were only two cases where there was substantial cross-resistance to multiple viruses—3R 49cM confers strong resistance to DMelSV and weak resistance to DObsSV, while 3L 46cM confers weak resistance to DMelSV and strong resistance to DObsSV.

Together, this modest number of loci with substantial effects on resistance explains most of the high genetic variance in resistance to the coevolved virus (Figure 4A, filled versus open circles). Individually resistant alleles cause an approximate 3-7 fold reduction in viral load (Figure 4C), and together they explain 59% of the genetic variance in susceptibility to DMelSV, 77% for DObsSV and 3% for DAffSV (Figure 4A, filled versus open circles). However, even after accounting for these genes there remains a significantly higher genetic variance in the viral load of the coevolved virus (Figure 4A, open circles, non-overlapping 95% CI).

## Discussion

We have found greater genetic variation in susceptibility to pathogens that naturally coevolve with a host compared to those that do not, suggesting that selection by pathogens acts to increase the amount of genetic variation in susceptibility. This effect was largely caused by a modest number of major-effect genes that explain over half of the genetic variance in resistance.

There are several different ways in which selection by pathogens could increase genetic variation. Negative frequency dependent selection can maintain genetic variation if the alleles conferring resistance are costly or increase susceptibility to other pathogen genotypes [18]. Directional selection can cause transient increases in genetic variance if it increases the frequency of rare or *de novo* resistance mutations [23]. Temporal or geographical variation in the prevalence of pathogens could maintain both resistant and susceptible alleles [20, 21].

As the genetic variants in *p62 (Ref(2)P)* and *CHKov1* that confer resistance to DMelSV have been identified, this has previously allowed us to use patterns of DNA sequence variation to infer how selection has acted on resistance in *D. melanogaster*. In both these genes the resistant alleles have arisen relatively recently by mutation and natural selection has pushed them rapidly up in frequency, leaving a characteristic signature of elevated linkage disequilibrium and low genetic diversity around the variant causing resistance [24, 25, 36]. There is no indication of negative frequency dependant selection, and these polymorphisms appear to have arisen from partial selective sweeps [24, 25].

At equilibrium directional selection on a trait is not expected to affect its genetic variance (relative to a population under mutation-drift balance; [17]). However, the genetic variance will transiently increase if the variants under selection are initially at low frequency [23], as was the case for both *p62* and *CHKov1* [24, 25]. A particular feature of pathogens is that the direction of selection is likely to continually change, as new pathogens appear in populations or existing pathogens evolve to overcome host defences. For example, in France and Germany in the 1980’s DMelSV evolved to largely overcome the effects of the resistant allele of *p62* [51, 52]. Similarly, DImmSV has swept through European populations of *D. immigrans* in the last ~16 years and DObsSV through UK populations of *D. obscura* in the last ~11 years [28, 31]. If selection by pathogens continually changes and resistance evolves from new mutations, then this may cause a sustained increase in genetic variance in susceptibility to infection.

Association studies in humans have revealed that quantitative traits are typically controlled by a very large number of genetic variants, each of which tends to have a very small effect [53, 54]. However, in humans some have advocated the view that susceptibility to infectious disease is qualitatively different from other traits and has a much simpler genetic basis [55]. Our results support this, with seven polymorphisms explaining over half the genetic variance. This confirms our previous work in *D. melanogaster* showing a simple genetic basis of virus resistance [11, 56]. As these genetic variants mostly only affect the naturally occurring pathogen of *D. melanogaster*, our results suggest that not only is selection by pathogens increasing the genetic variance but it is also altering the genetic architecture of the trait by introducing major-effect variants into the population. One explanation for this observation is that most quantitative traits are under stabilising selection, so major effect variants will tend to be deleterious and removed by selection [57]. In contrast, selection by pathogens likely changes through time and populations may be far from their optimal level of resistance. If this is the case, Fisher’s geometric model predicts that major effect variants will be favoured by directional selection [58]. Similar arguments may apply if resistance polymorphisms are under negative frequency dependant selection.

A major source of emerging infectious disease is pathogens jumping into novel hosts where they have no co-evolutionary history [59, 60]. Our results suggest that when a pathogen infects a novel host species, there may be far less genetic variation in susceptibility among individuals than is normally the case. This may create a ‘monoculture effect’ [6, 61, 62], which could leave populations vulnerable to epidemics of pathogens that have previously circulated in other host species. Longer term, low levels of pre-standing genetic variation may also slow down the rate at which the new host can evolve resistance to a new pathogen.

In conclusion, we have demonstrated that selection by pathogens can increase the amount of genetic variation in host susceptibility and result in a simple genetic architecture where resistance is largely controlled by major effect resistance loci.

## Methods

### Virus extraction and infection

We extracted the sigma viruses DAffSV, DImmSV, DMelSV and DObsSV from infected stocks of *D. affinis* (line: NC10), *D. immigrans* (line: DA2), *D. melanogaster* (line: E320 Ex) and *D. obscura* (line: 10A) respectively [28-31, 44]. These infected lines were collected from the wild between 2007-2012 (all from the UK, bar *D.affinis* which was collected in the USA). Infected fly stocks were checked for infection by exposing the flies to 100% CO_2_ at 12°C for 15mins then paralysed flies were collected 30mins later. The DImmSV infected line does not show CO_2_ sensitivity and so was confirmed to have a high level of infection using RT-PCR. Infected flies were frozen at −80°C and later homogenised in ringers solution (2.5μl per fly) and centrifuged at 13,000g for 10min at 4°C. The supernatant was collected, 2% v/v FBS was added then virus solutions were aliquoted and stored at −80°C.

### Experimental design

We set up a common garden experiment to measure genetic variation in susceptibility to natural and non-natural viruses across four host species. For each species we infected it with its own virus, as well as two viruses that do not infect that host species (see Figure 1). All fly stocks used were tested for existing sigma virus infection using RT-PCR over two generations. For all species we collected flies from the wild and we used a full-sib mating design. The progeny of these crosses was infected by injecting them with the viruses intrathoracically and measuring viral RNA loads 15 days post infection, as in [63]. This time point was selected as RNA viral load tends to plateau from around day 15 post infection and there is no significant mortality from infection in this period. The specifics for each host species are detailed in the supplementary methods.

### Known resistance genes in *D. melanogaster*

We genotyped parents of each *D. melanogaster* full sib family from the experiment above for two resistance alleles that are known to confer protection against DMelSV; *p62* (*Ref(2)P)* and *CHKov1*. We genotyped parental flies using PCR assays that produce different sized products depending on whether flies carry resistant or susceptible alleles. Information on these PCRs and primer sequences can be found in table S3. We then calculated the number of resistance alleles in each family by summing the number of alleles from both mothers and fathers. We produced genotype information for 230 of the 255 families.

Another resistance allele has been identified in the gene *Ge-1* [34]. However, this allele has been found to occur at a low frequency in wild populations. We genotyped a subset of 184 parental flies from our experiment and found the resistant allele was not present, suggesting it is rare or absent.

We further examined the effect of alleles known to affect susceptibility to DMelSV on all three viruses. Firstly, we infected 32 lines from the Drosophila Genetic Reference Panel (DGRP) [49] that were susceptible for both *p62* and *CHKov1* (*n*=11 lines), were resistant for *CHKov1* only (*n*=13 lines), or were resistant for both genes (*n*=8 lines) with DMelSV, DAffSV and DObsSV. No lines in the panel were resistant for *p62* and susceptible for *CHKov1*. We infected a mean of 18 flies per line (range = 3-22).

### Mapping resistance genes in *D. melanogaster*

We used 377 DSPR lines (154 from panel A and 223 from panel B, http://FlyRILs.org [50, 64], kindly provided by S.J. Macdonald, University of Kansas) to carry out a Quantitative Trait Locus (QTL) study to examine the genetic basis of resistance to DAffSV, DMelSV and DObsSV in *D. melanogaster*.

3 females and 3 males from each DSPR line were placed into yeasted cornmeal vials and allowed to lay for 3-4 days, at 25°C. Male offspring were collected at 0-4 days post-eclosion, and placed at 18°C for 4-6 days. Flies were then injected with DMelSV, DAffSV or DObsSV as described above. Injected males were maintained on unyeasted cornmeal at 18°C, and frozen on day 15 post-infection as above.

Injections were carried out over 13 weeks. Each day of injection a mean of 47 unique lines (range 20-60) and 51 replicate vials were injected with 1-3 different viruses. In total we assayed 377 DSPR lines (108 lines had 2 biological replicates). Each replicate vial contained a mean of 12 flies (range 1-22). In total, 15,916 flies were injected across both panels of DSPR lines. We injected 319 lines with all 3 viruses, 38 with 2 viruses and 20 with 1 virus. The order of injection of lines and of viruses, was randomised across injection days. Independent biological replicates were injected on different days. Panel A and Panel B lines were assayed in two overlapping blocks.

### Measuring viral load

We measured the change in RNA viral load using qRT-PCR. The viral RNA load was expressed relative to the endogenous control housekeeping gene *RpL32* (*Rp49*). RNA was extracted from flies homogenised in Trizol and reverse transcribed with GoScript reverse transcriptase (Promega) and random hexamer primers, and then diluted 1:10 with nuclease free water. The qRT-PCR was performed on an Applied Biosystems StepOnePlus system using Sensifast Hi-Rox Sybr kit (Bioline) with the following PCR cycle: 95°C for 2min followed by 40 cycles of: 95°C for 5 sec followed by 60°C for 30 sec. Two qRT-PCR reactions (technical replicates) were carried out per sample with both the viral and endogenous control primers. Each qRT-PCR plate contained three standard samples, and all experimental samples were split across plates in a blocked design. A linear model was used to correct the cycle threshold (Ct) values for differences between qRT-PCR plates. Primer sequences are in Table S6.

To estimate viral load, we calculated ΔCt as the difference between the qRT-PCR cycle thresholds of the virus and the endogenous control. Viral load calculated without using the endogenous control is strongly correlated to ΔCt for all species.

### Statistical analysis full-sib experiments

We used a linear mixed model to examine the amount of genetic variation in susceptibility to the different viruses. We used a trivariate model with the load of the three viruses as the response variable. For each species the model was structured as:

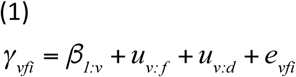

Where *y*_*vfi*_ is the log_2_ viral load of the *i^th^* biological replicate of full-sib family *f* infected with virus *v. β* are the fixed effects, with *β*_1_ being the mean viral load of each virus. *u* are the random effects for full-sib families (*f*) and for the day of injection (*d*), *e* are the residuals. By assuming that all the genetic variation in the population is additive [65], we estimated the genetic variance (*V_G_*) of the viral load as twice the between-family variance [14, 15]. Both empirical data and theory suggest additive genetic variation makes up large proportion of the total genetic variance [65].

In addition, for *D. melanogaster* we ran a further model that included the additional fixed effects *β*_*2:v*_ and *β*_*3:v*_ that are the linear effects of the *CHKov1* and *p62* (*Ref(2)P)* resistance alleles. We assumed these genes had additive effects, and modelled their effects simply as the proportion of resistant alleles in a family (if one parent was heterozygous and the other homozygous susceptible, the value is 0.25).

The model was fitted using the MCMCglmm package in R [66]. The random effects (and residuals) are assumed to be multivariate normal with zero mean and covariance structure **V**⊗ **I. I** is an identity matrix, and **V** a matrix of estimated variances and covariances. For the random effects **V** is a 3×3 covariance matrix describing the variances for each virus and the covariances between them. The off-diagonal elements of **V** for the residual were set to zero because the covariances between traits at these levels are not estimable by design.

Diffuse independent normal priors were placed on the fixed effects (means of zero and variances of 10^8^). Parameter expanded priors were placed on the covariance matrices resulting in scaled multivariate *F* distributions which have the property that the marginal distributions for the variances are scaled (by 1000) *F*_1, 1_. The exceptions were the residual variances for which an inverse-gamma prior was used with shape and scale equal to 0.001. The MCMC chain was ran for 130 million iterations with a burn-in of 30 million iterations and a thinning interval of 100,000.

We confirmed the results were not sensitive to the choice of prior by also fitting models with inverse-Wishart and flat priors for the variance covariance matrices (described in [63]), as well as fitting the models by REML in ASReml R [67]. These analyses all gave qualitatively similar results (data not shown).

### Statistical analysis of QTL experiment

Based on genotyping data, the probability that each Recombinant Inbred Line (RIL) in the DSPR panel was derived from each of the 8 founder lines has been estimated at 10kB intervals across the genome (King et al. 2012). To identify QTL affecting viral load we first calculated the mean viral load (ΔCt) across the biological replicates of each RIL. We then regressed the mean viral load against the 8 genotype probabilities and calculated logarithm of odds (LOD) scores using the DSPRqtl package in R (King et al. 2012). These LOD scores were calculated separately for DSPR Panel A and Panel B, and then summed at each genomic location. To obtain a significance threshold, we permuted our mean viral load estimates across the RILs within each panel, repeated the analysis above and recorded the highest LOD score across the entire genome. This process was repeated 1000 times to obtain a null distribution of the maximum LOD score.

To estimate the effect of each QTL we assumed that there was a single genetic variant affecting viral load, so the founder alleles could be assigned to two functionally distinct allelic classes. First, we regressed the mean viral load against the genotype probabilities (as described above), resulting in estimates of the mean viral load of each founder allele in the dataset. The two DSPR panels had one founder line (line 8) in common. For QTL where the line 8 allele was present in both panels, this analysis included data from both panels, and ‘panel’ was included as a fixed effect in the analysis. When this was not the case, we analysed only data from the panel where the QTL was most significant. We then ranked the founder alleles by viral load estimate, and split this ranked list into all possible groups of two alleles. For each split the genotype probabilities in the first group of founder alleles were summed. We then regressed mean viral load against each of these combined genotype probabilities. The regression model with the highest likelihood was taken as the most likely classification into allelic classes. The effect size of the QTL was then estimated from this model.

To estimate the genetic variance in viral load within the DSPR panels we modified the model described in Equation 1 as follows. *γ*_*vfi*_ is the log_2_ viral load of the *i^th^* biological replicate of each RIL *f* infected with virus *v.* There was a single fixed effect, *β*, of the panel the line is from. *u* are the random effects for each RIL (*f*). The day of injection (*d*) was omitted. As all the RILs are homozygous, we estimated the genetic variance in viral load (*V_G_*) as half the between-RIL variance. This assumes all the genetic variation is additive.

To estimate the proportion of the genetic variance that is explained by the QTL we identified, we repeated this analysis but included the 7 QTL we identified as fixed effects in the model. Each QTL was included by estimating the probability that each line carried the resistant allele of the QTL and adding this as a fixed effect to the model. The between-RIL variance then allowed us to estimate the genetic variance in viral load after removing the effects of the QTL.

## Data availability

Datasets and R code for estimating the amount of genetic variation in susceptibility https://doi.org/10.6084/m9.figshare.6743339DGRP dataset https://doi.org/10.6084/m9.figshare.6743354 DSPR dataset and R code https://doi.org/10.6084/m9.figshare.7195751

## Acknowledgements

Many thanks to: Alastair Wilson and Jarrod Hadfield for useful advice and discussion; Stuart Macdonald for providing DSPR lines, Trudy Mackay for providing the DGRP fly lines and Kelly Dyer with help collecting *D.affinis*; Darren Obbard for providing photographs of Drosophila species; Camille Bonneaud, Katherine Roberts, Ryan Imrie and the Unckless lab group for constructive comments on this work.

## Supplementary materials

**Figure S1.**
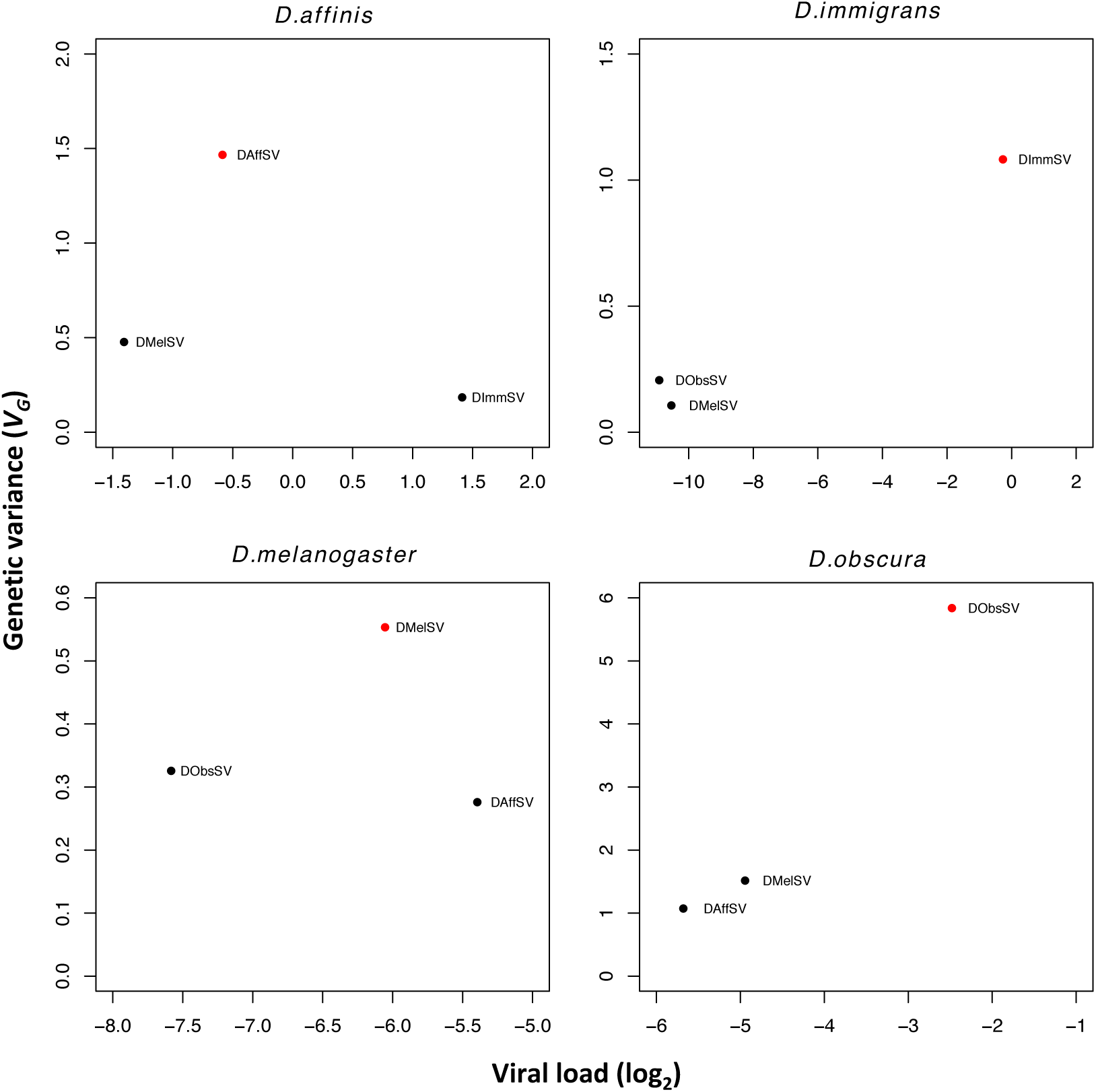
Estimates of genetic variance plotted against mean viral load for each species-virus.

**Table S1.** Credible intervals of the differences between estimates of genetic variation in susceptibility across host species and viruses. The natural virus for each host is in red and bold. 95% CIs show differences in estimates of genetic variation for different host-virus combinations, intervals that do not cross zero represent statistically significant differences.

| Host | Virus comparison | 95% CIs |  |
| --- | --- | --- | --- |
| *D. affinis* | **DAffSV**-DImmSV | 0.862 | 1.699 |
|  | **DAffSV**-DMelSV | 0.532 | 1.510 |
|  | DImmSV-DMelSV | -0.606 | 0.004 |
| *D. immigrans* | **DImmSV**-DMelSV | 0.630 | 1.331 |
|  | **DImmSV**-DObsSV | 0.500 | 1.240 |
|  | DMelSV-DObsSV | -0.365 | 0.167 |
| *D. melanogaster* | **DMelSV**-DAffSV | 0.080 | 0.485 |
|  | **DMelSV**-DObsSV | 0.012 | 0.486 |
|  | DAffSV-DObsSV | -0.264 | 0.149 |
| *D. obscura* | **DObsSV**-DAffSV | 3.219 | 6.184 |
|  | **DObsSV**-DMelSV | 2.786 | 5.817 |
|  | DAffSV-DMelSV | -0.996 | 0.102 |

**Table S2.** Estimates of genetic variation for each host virus combination. The natural virus for each host is in red and bold.

| Host | Virus | Mean | 95% CIs |  |
| --- | --- | --- | --- | --- |
| *D.affinis* | **DAffSV** | 1.47 | 1.12 | 1.92 |
| *D.affinis* | DImmSV | 0.18 | 0.05 | 0.32 |
| *D.affinis* | DMelSV | 0.48 | 0.21 | 0.77 |
| *D.immigrans* | **DImmSV** | 1.08 | 0.81 | 1.39 |
| *D.immigrans* | DMelSV | 0.11 | 0.00 | 0.27 |
| *D.immigrans* | DObsSV | 0.21 | 0.00 | 0.41 |
| *D.melanogaster* | **DMelSV** | 0.55 | 0.39 | 0.73 |
| *D.melanogaster* | DAffSV | 0.28 | 0.16 | 0.39 |
| *D.melanogaster* | DObsSV | 0.33 | 0.15 | 0.48 |
| *D.obscura* | **DObsSV** | 5.84 | 4.39 | 7.23 |
| *D.obscura* | DAffSV | 1.07 | 0.75 | 1.44 |
| *D.obscura* | DMelSV | 1.52 | 1.01 | 1.98 |

**Table S3.** Genetic correlations (*r_g_*) in viral loads across host species after infection by coevolved and non-coevolved viruses. The coevolved (natural) virus for each host is in red and bold. Models ran using REML gave similar estimates.

| Host | Viruses | Mean | 95% CI |
| --- | --- | --- | --- |
| *D. affinis* | **DAffSV**-DImmSV | 0.50 | -0.01, 0.92 |
|  | **DAffSV**-DMelSV | 0.05 | -0.54, 0.52 |
|  | DMelSV-DImmSV | 0.02 | -0.62, 0.57 |
| *D. immigrans* | **DImmSV**-DMelSV | -0.32 | -0.91, 0.33 |
|  | **DImmSV**-DObsSV | -0.22 | -0.70, 0.26 |
|  | DMelSV-DObsSV | 0.29 | -0.43, 0.94 |
| *D. melanogaster* | **DMelSV**-DAffSV | 0.40 | 0.20, 0.61 |
|  | **DMelSV**-DObsSV | 0.25 | -0.02, 0.48 |
|  | DAffSV-DObsSV | 0.36 | 0.10, 0.62 |
| *D. obscura* | **DObsSV**-DAffSV | 0.21 | -0.03, 0.46 |
|  | **DObsSV**-DMelSV | 0.40 | 0.16, 0.63 |
|  | DAffSV-DMelSV | 0.68 | 0.49, 0.84 |

**Table S4.**
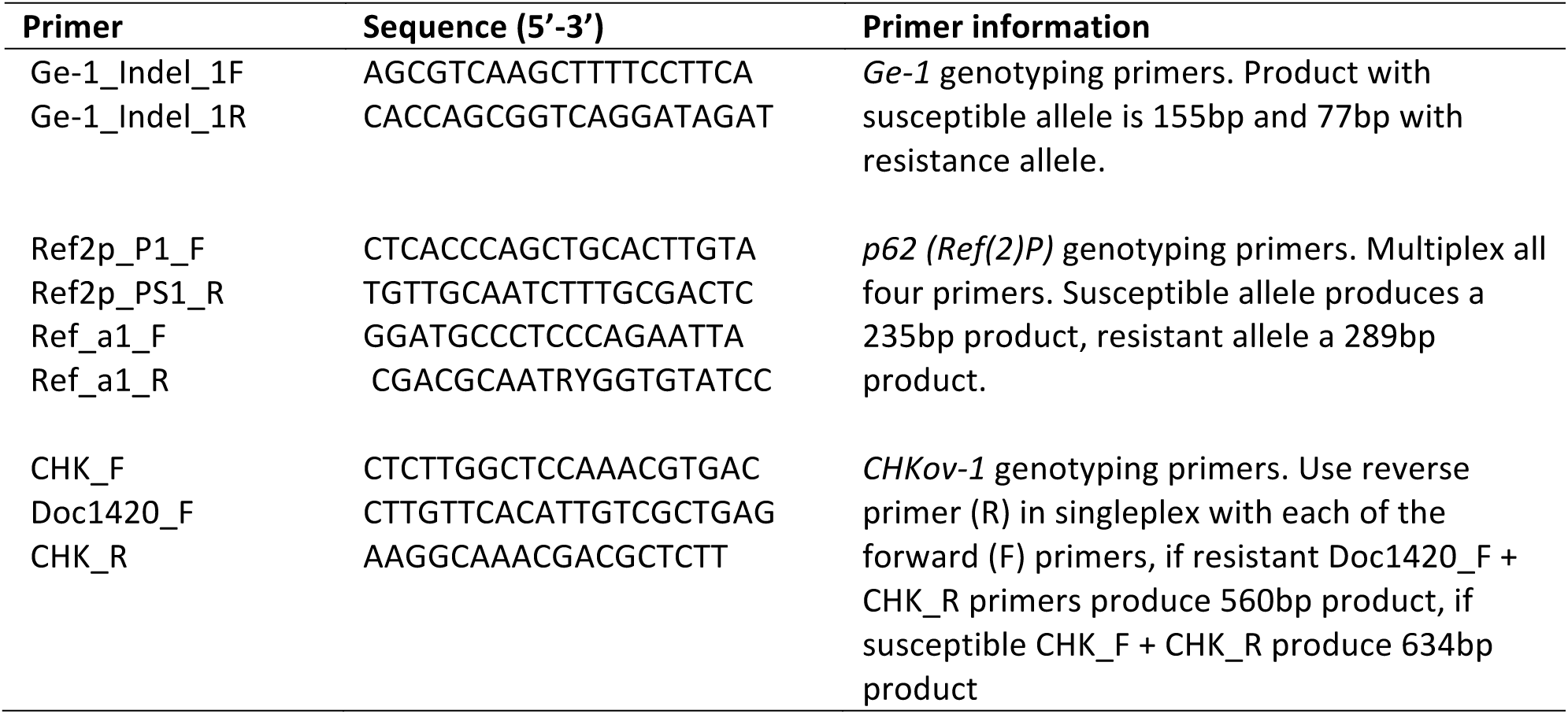
Primers for genotyping *D. melanogaster* resistance genes *Ge-1, p62 (Ref(2)P) and CHKov-1.* PCRs were carried out using a touchdown PCR cycle (95°C 30sec, 62°C (−1°C per cycle) 30sec, 72°C 1min; for 10x cycles followed by; 95°C 30sec, 52°C 30sec, 72°C 1min; for a further 25x cycles).

**Table S5.**
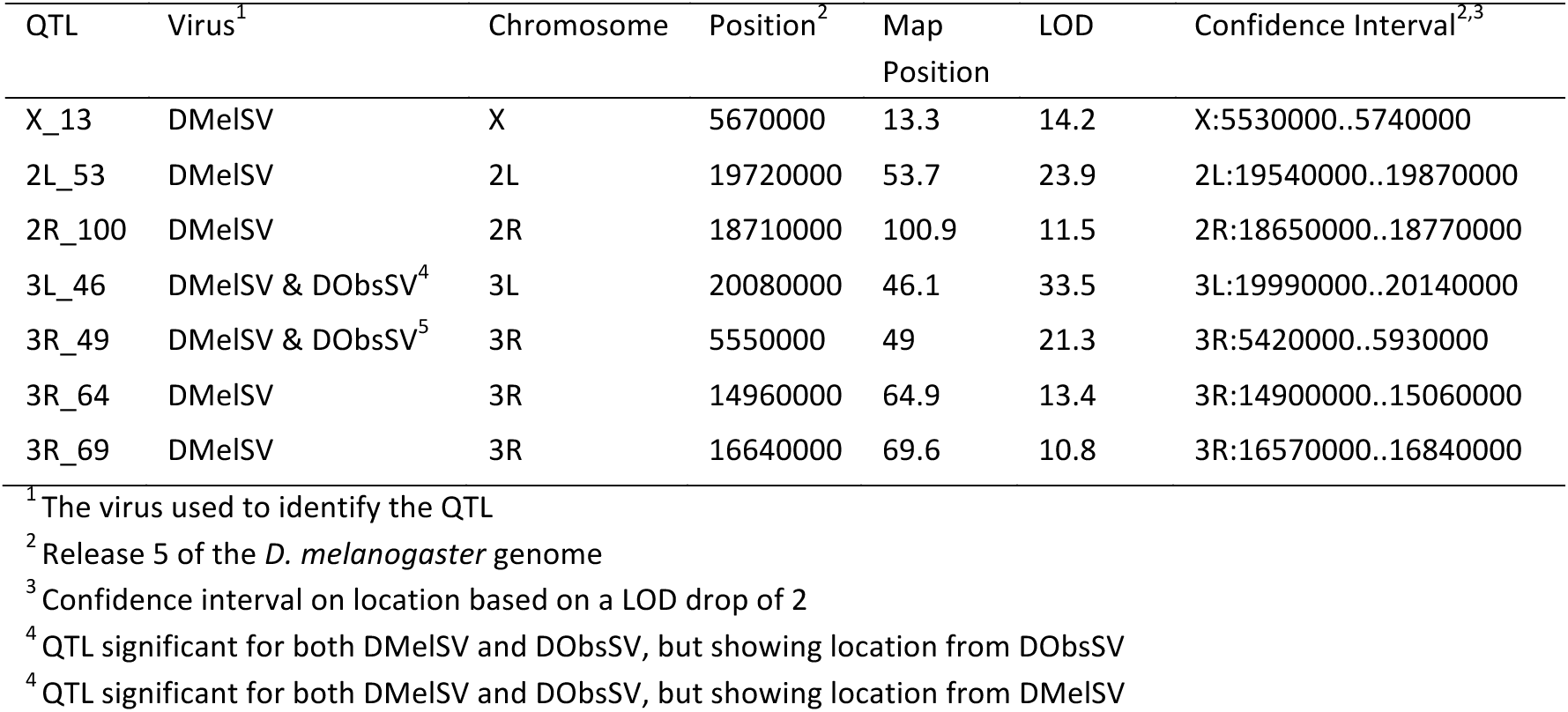
QTL and their locations.

**Table S6.**
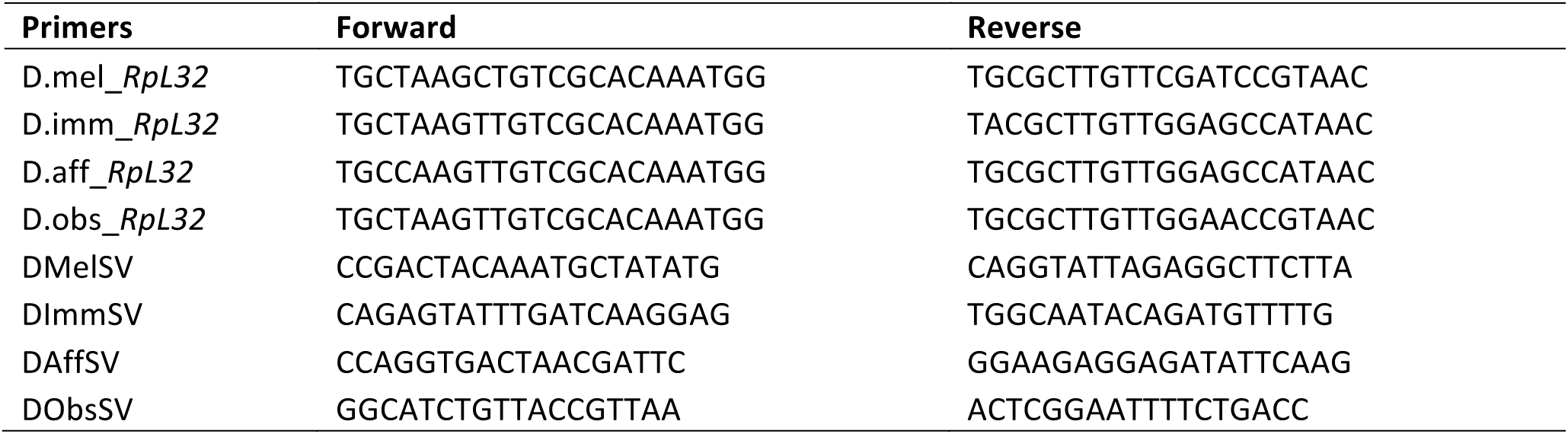
Primers for qRT-PCR (5’-3’).

## Supplementary Methods

All lines were screened for their retrospective sigma virus over two generations by RT-PCR, and infected isofemale lines discarded prior to the experiment.

### Drosophila melanogaster

We created an outcrossed population by combining 150 isofemale lines of *D. melanogaster* (collected in Accra, Ghana (5.593, −0.188) in 2014) in a population cage. The population was maintained throughout the experiment with a large population size (~1500-2000 flies), with eggs collected from the population cage used to set up each subsequent generation. All rearing was carried out on cornmeal medium (recipe below) sprinkled with live yeast (‘yeasted’) at 25°C.

Virgin flies were collected daily from bottles set up at a controlled egg density. Full-sib families were set up using crosses of single male and female virgins placed in the same vial and aged for 3 days. Each of these families was tipped onto fresh food daily for 5 days to create replicate vials. After 5 days the adult flies were frozen for later genotyping. 12 days after laying, male offspring were collected from each replicate vial and split into two vials of cornmeal medium without any yeast on the surface (‘unyeasted’) and placed at 18°C. After 5 days these flies were injected with 69nl of virus extract intra-abdominally using a Nanoject II micro-injector (Drummond scientific). Injected flies were kept at 18°C and tipped onto fresh unyeasted cornmeal every 5 days, before being homogenised in Trizol (Invitrogen) and frozen at −80°C on day 15 post injection for later RNA extraction and qRT-PCR.

Injections were carried out over 25 overlapping blocks. Each block consisted of 10 families, and each day, 2 vials per family were injected with 2 different viruses. Each replicate vial contained a mean of 14 flies (range 3-28 flies). In total we measured 255 families over 1567 biological replicates. We aimed to carry out a minimum of 2 replicates of each virus per family, but where possible we carried out 3 or 4 replicates (92 virus-family combinations had 1 replicate, 555 had 2 replicates, 82 had 3 replicates and 25 had 4 replicates.). 248 families had replicates for all 3 viruses. Blocks were staggered to overlap with at least 20 families being infected on any one day. The order families were injected in was randomised, and the order the different viruses were injected was blocked across days.

### D. immigrans

92 *D. immigrans* lines were collected from Madingley, Cambridge, UK (52.225, 0.043) in 2012 and 2015. Full-sib families were set up using crosses between the 92 isofemale lines of *Drosophila immigrans*. Flies were reared on malt food (recipe below) at 18°C. Crosses were between different isofemale lines (i.e. excluding reciprocal crosses) and maximising the number of lines used. Families were established from 2-4 day old single female and male virgin flies placed in the same vial for 7 days. These crosses were tipped onto fresh food every 7 days to generate replicate vials of each family. Eclosed males were collected 27-34 days after initial egg laying and injected with DImmSV, DMelSV or DObsSV 1-3 days post-collection, then maintained and frozen on day 15 post-infection as above.

Injections were carried out over 18 overlapping blocks. Each block consisted of an average of 19 families and 46 replicate vials. Each replicate vial contained a mean of 14 flies (range: 4-26). In total we assayed 341 families over 812 biological replicates. We aimed to have a minimum of 2 replicates per virus per family (235 virus-family combinations had 1 replicate, 270 had 2 replicates, 11 had 3 replicates and 1 had 4 replicates). 140 families had replicates across 2 different viruses and 18 families had replicates for all 3 viruses. Blocks were staggered to overlap with a mean of 39 families being infected on any one day. The order families were injected in was randomised, and the order the different viruses were injected was blocked across days.

### D. affinis

| Site | N |
| --- | --- |
| Athens, Georgia, USA, (33.946, -83.384) in 2012 | 13 |
| Great Smokey Mountain National Park, Gatlinburg, USA (35.698, -83.613) in 2015 | 23 |
| Rochester, New York, USA (43.135, -77.599) in 2012 | 4 |

Full-sib families were set up using crosses between 40 isofemale lines of *Drosophila affinis* (see above for collection details) collected in the U.S. Flies were reared on malt food (recipe below) at 18°C. Crosses were between different isofemale lines (i.e. excluding reciprocal crosses) and maximising the number of lines used. Families were established from 6 day old single female and male virgin flies placed in the same vial for 7 days. These crosses were tipped onto fresh food every 7 days to generate replicate vials of each family. Eclosed males were collected 35-42 days after initial egg laying and then injected with DAffSV, DImmSV or DMelSV 1-3 days post-collection, then maintained and frozen on day 15 post-infection as above.

Injections were carried out over 27 overlapping blocks. Each block consisted of an average of 19 families and 28 replicate vials. Each replicate vial contained a mean of 11 flies (range: 3-23). In total we assayed 520 families over 1003 biological replicates. We aimed to have a minimum of 2 replicates per virus per family (336 virus-family combinations had 1 replicate, 286 had 2 replicates and 30 had 3 replicates). 109 families had replicates across 2 different viruses and 12 families had replicates for all 3 viruses. Blocks were staggered to overlap with a mean of 23 families being infected on any one day. The order families were injected in was randomised, and the order the different viruses were injected was blocked across days.

### D. obscura

| Site | N |
| --- | --- |
| Derbyshire Site A , UK (52.978, -1.440) in 2012 | 4 |
| Derbyshire Site C, UK (52.903, -1.374), in 2012 | 1 |
| Les Gorges du Chambon, France (45.622, 0.555) in 2012 | 1 |
| Madingley, Cambridge, UK (52.226, 0.046) in 2014 | 15 |

D. *obscura* were collected in the United Kingdom and France (see above). Males and females were separated, and females were placed in vials to establish isofemale lines. Full-sib families were set up using crosses between 21 isofemale lines of *Drosophila obscura* collected in the UK. Flies were reared on banana food (recipe below) at 18°C. Crosses were between different isofemale lines (i.e. excluding reciprocal crosses) and maximising the number of lines used. Families were established from 6 day old single female and male virgin flies placed in the same vial for 7 days. These crosses were tipped onto fresh food every 7 days to generate replicate vials of each family. Eclosed males were collected 35-42 days after initial egg laying and then injected with DAffSV, DMelSV, or DObsSV 1-3 days post-collection, then maintained and frozen on day 15 post-infection as above.

Injections were carried out over 25 overlapping blocks. Each block consisted of a mean of 16 families with a mean of 76 vials being injected each day for 12 days. Each replicate vial contained a mean of 8 flies (range: 1-15). In total we assayed 320 families over 913 biological replicates. We aimed to have a minimum of 2 replicates per virus per family (126 virus-family combinations had 1 replicate, 314 had 2 replicates and 49 had 3 replicates and 3 had 4 replicates). 94 families had replicates across 2 different viruses and 39 families had replicates for all 3 viruses. Blocks were staggered to overlap with at least 10 families being infected on any one day. The order families were injected in was randomised, and the order the different viruses were injected was blocked across days.

### Sample size estimation

The number of full-sib families required for estimating genetic variance in susceptibility was determined by simulation using previous estimates of genetic variation to DMelSV in *D. melanogaster* [11]. After carrying out the full-sib experiment in *D. melanogaster*, we then down-sampled this data to calculate the minimum number of families required to provide accurate estimates for the other species. Sample sizes for the DSPR experiment were based on previous data [56].

### Food recipes

Banana:

Mixture 1 1000ml water 30g yeast

10g agar

Mixture 2 20ml Nipagin

150g pureed banana 50g corn syrup

30g malt powder

Bring mixture 1 to the boil for 3-4 minutes, whisk constantly. Add to Mixture 2 to Mixture 1. Whisk constantly and simmer for 5 minutes.

Cornmeal:

1200ml water 13g agar

105g dextrose 105g maize 23g yeast

Combine and bring to a boil for 5mins, cool to 70°C before adding 35ml Nipagin (10%)

Malt:

1000ml water 10g agar

60g semolina 20g yeast

80g malt extract

Combine and bring to a boil for 5mins, cool to 70°C and then add 14ml Nipagin (10%) and 5ml proprionic acid.

